# Ultra-Slow Rhythmic Movement in the Elongating Tail of *Xenopus* Tadpoles

**DOI:** 10.64898/2026.05.27.728187

**Authors:** Iroha Nakane, Yuki Asakura, Soichiro Kato

## Abstract

Since Darwin’s time, ultra-slow oscillatory movements have been recognized as a characteristic feature of axis elongation in plant organs such as shoots and tendrils, yet have not been reported in elongating animal body axes. Here, using long-term imaging of multiple anesthetized *Xenopus laevis* embryos with a modified document scanner, we identify a previously unrecognized ultra-slow rhythmic movement in the developing tadpole tail. This movement emerged during tailbud development, persisted once established, and occurred at an ultra-low frequency of approximately 10^−4^ Hz, several orders of magnitude slower than known active animal tail movements. Temperature-dependent imaging showed that its period varied with developmental rate and converged after time rescaling based on tail elongation rate, indicating a close link to morphogenetic elongation rather than conventional locomotor behavior. The timescale and physical context of this movement resemble plant circumnutation, in which slow organ-scale movement arises during constrained elongation. These findings suggest that slow rhythmic motion may represent a morphogenetic mode of animal movement and may be a recurrent dynamic feature of constrained elongation in living structures.

## Introduction

Charles Darwin and his son Francis described circumnutation in 1880 as an oscillatory movement of growing plant organs such as stems and tendrils, unfolding over timescales of hours to days (1). In *The Power of Movement in Plants*, they argued that these slow movements are intrinsic to plant growth and contribute to adaptive interactions with the environment. More recently, circumnutation has been interpreted as arising from endogenous oscillatory activity coupled with feedback responses (2), and as helping organs avoid obstacles and explore spaces more favorable for growth (3).

Rhythmic dynamics also accompany tissue elongation in animals, yet these rhythms are generally viewed as internal patterning or force-generating processes rather than as slow, organ-scale movements. For example, the elongating vertebrate body axis contains an intrinsic temporal rhythm: the segmentation clock periodically patterns the paraxial mesoderm into somites (4). In other systems, rhythmic force-generating processes contribute more directly to elongation, including cyclic muscle contractions in *C. elegans* embryos (5) and pulsatile actomyosin contractions during *Drosophila* egg chamber elongation (6), germ-band extension (7), and *Xenopus* convergent extension (8). However, no comparable slow, organ-scale oscillatory movement has been described in an elongating animal body axis.

Here, we report an ultra-slow periodic tail movement in developing *Xenopus laevis* tadpoles that emerges during tail elongation and whose period scales with the rate of tail elongation. These findings suggest that slow rhythmic movement may be a shared dynamic feature of tissue elongation across plants and animals.

## Results and Discussion

We developed a custom long-term imaging system based on a modified document scanner to monitor multiple anesthetized embryos simultaneously for more than two days while avoiding repeated repositioning. Using this system, we identified slow rhythmic left–right oscillations of the tail tip during mid-tailbud development (Fig. 1A–D, Video S1). To isolate this oscillatory component from the slower trend associated with tail elongation, we detrended the tail-tip trajectory using locally estimated scatterplot smoothing (LOESS) (Fig. 1E). Wavelet analysis of the detrended signal revealed a clear periodic component around 100 min that, once it emerged, generally persisted with little change in period until at least St. 41 (Fig. 1F). To compare onset across individuals, we extracted the peak-period band (±50%) from each scalogram and aligned these across 15 embryos, showing that periodicity emerged across individuals between approximately St. 30 and St. 40 (Fig. 1G). The oscillation period was on the order of hours, corresponding to an ultra-low frequency of approximately 10^−4^ Hz (Fig. 1E, F). This frequency is several orders of magnitude lower than the 10–25 Hz tail beating used for swimming in *Xenopus* tadpoles, making it unlikely to represent a simple variant of previously described locomotor tail movements (9). More broadly, reported frequencies of active animal tail movements range from ∼0.15 Hz during Greenland shark swimming to 60–100 Hz during rattlesnake sound production, still far above the frequency observed here (10,11).

**Figure 1.**
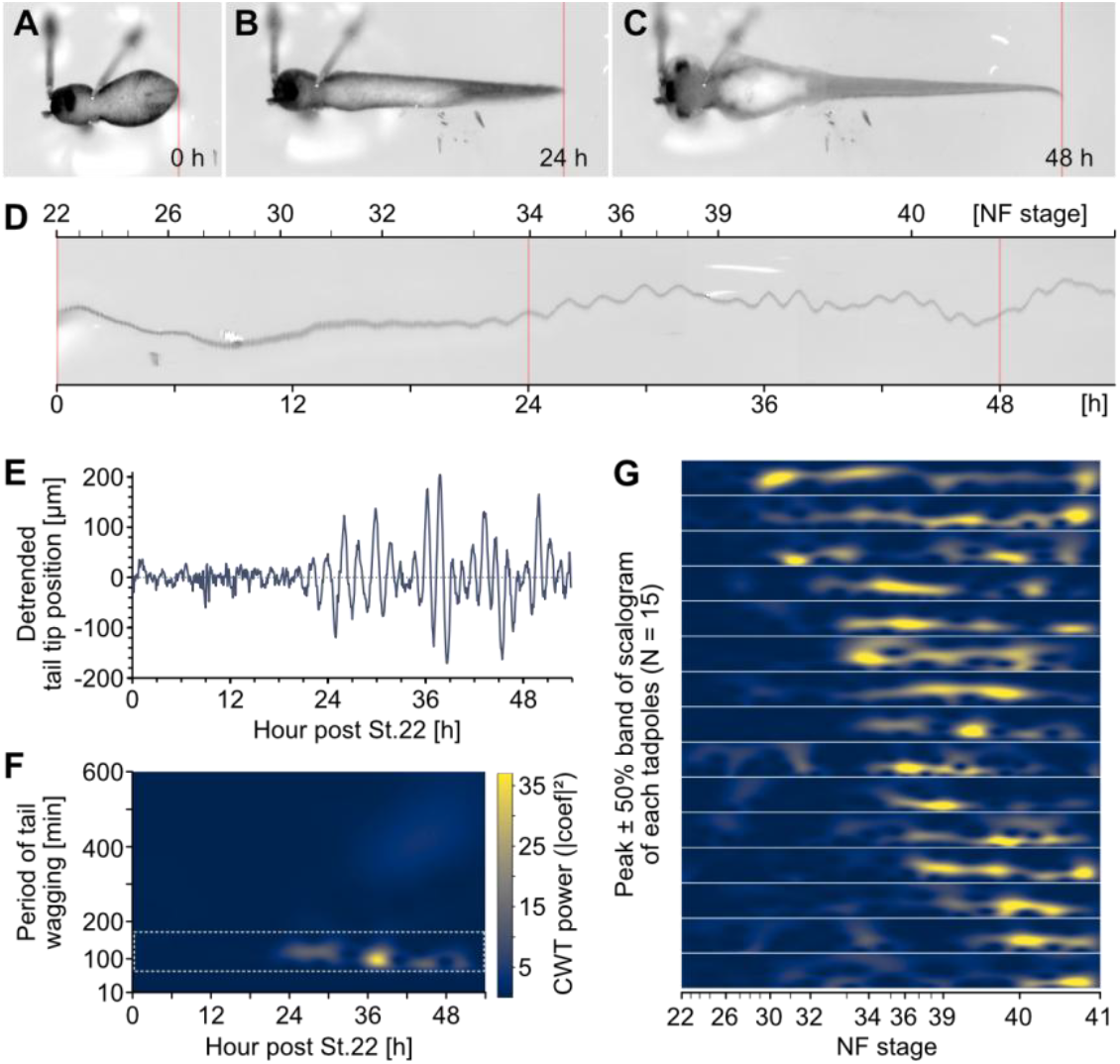
Ultra-slow rhythmic movement emerges in the elongating tail of Xenopus tadpoles. (A–C) Representative ventral images of the same anesthetized embryo at 0, 24, and 48 h during long-term time-lapse imaging starting at St. 22. Red bands indicate the tail-tip region used to generate the temporal montage in (D). (D–F) Analysis of the same embryo shown in (A–C). (D) Temporal montage of the tail-tip region. (E) LOESS-detrended lateral displacement of the tail tip. (F) Wavelet power scalogram corresponding to the detrended trajectory shown in (E). (G) Peak-period bands (±50% of the peak period) extracted from wavelet scalograms for 15 embryos and aligned by Nieuwkoop–Faber stage (St. 22–41). The mean peak period was 145.9 ± 34.1 min (mean ± SD; n = 15 embryos from 3 independent clutches).

Because ultra-slow movement occurs on a timescale closer to morphogenesis than to swimming behavior, we asked whether its period scales with developmental rate. To test this, we maintained embryos at 13°C, 18°C, or 23°C during long-term imaging, thereby altering developmental rate by approximately threefold. We then quantified detrended tail-tip motion under each temperature condition (Fig. 2A). Over the imaging interval, tail elongation was nearly linear, with the elongation rate differing across temperature conditions (Fig. 2B). The oscillation period also varied with developmental rate, being longest at 13°C and shortest at 23°C (Fig. 2A, Video S2). Fast Fourier transform (FFT)-based analysis confirmed this trend, yielding mean ± SD periods of 272.0 ± 40.9 min at 13°C, 167.7 ± 10.9 min at 18°C, and 110.4 ± 9.6 min at 23°C (Fig. 2C).

**Figure 2.**
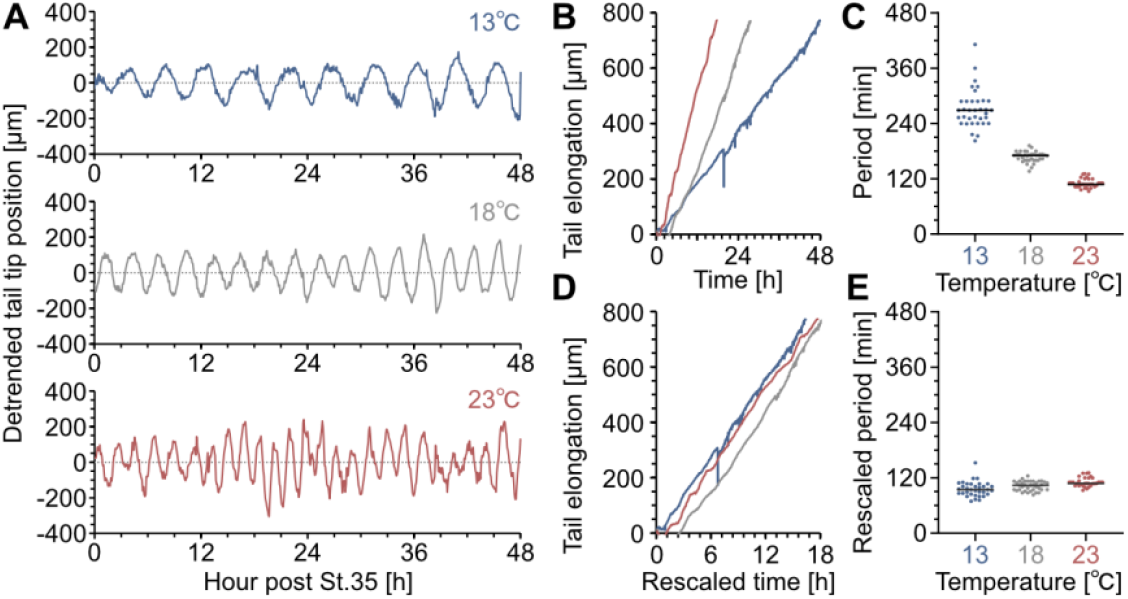
Developmental rescaling aligns the periods of rhythmic tail movement across temperatures. (A) Representative LOESS-detrended tail-tip displacement traces over 48 h from NF stage 35 onward at 13°C (blue, top), 18°C (gray, middle), and 23°C (red, bottom). (B) Representative tail elongation trajectories before time rescaling. (C) FFT-estimated oscillation periods before rescaling. (D) Representative tail elongation trajectories after time rescaling based on fitted tail elongation slopes. (E) FFT-estimated oscillation periods after rescaling to the fitted tail elongation rate at 23°C. Dots indicate individual embryos; horizontal bars indicate means. n = 35, 38, and 32 embryos for 13°C, 18°C, and 23°C, respectively, pooled from 4 independent clutches, each represented in all three temperature conditions.

To test whether ultra-slow movement scales with tail elongation, we rescaled the time axis in each temperature condition using the slope of a linear fit to tail elongation, so that the fitted elongation rate in each condition matched that at 23°C. After rescaling, the fitted elongation trajectories were closely aligned across temperatures (Fig. 2D), and the oscillation periods converged to similar mean ± SD values: 95.5 ± 13.7 min at 13°C, 102.4 ± 10.8 min at 18°C, and 110.7 ± 9.6 min at 23°C (Fig. 2E, Video S3). Together, these results suggest that the ultra-slow tail oscillation is linked to morphogenetic elongation.

Although the origin of this ultra-slow periodicity remains unknown, elongating plant tissues provide a useful physical point of comparison. Slow rhythmic movements such as circumnutation are well documented in growing plant structures, including shoots and tendrils, and occur over timescales ranging from hours to days (1–3). The ultra-slow movement observed in *Xenopus* tadpoles falls within this range. In both systems, elongation involves pressure-driven expansion under circumferential structural constraint: in plants, turgor-driven expansion acts against cellulose-rich cell walls whose anisotropy is linked to cortical microtubule orientation (12), whereas in *Xenopus*, vacuolization-driven expansion of notochord cells occurs within a collagenous sheath, contributing to axial elongation and body straightening (13, 14). Thus, despite profound differences in molecular composition and body plan, both systems represent elongating structures in which internal expansion is constrained by anisotropic external or circumferential architecture.

Plant circumnutation is thought to arise when spatially and temporally varying differential growth, shaped by organ mechanics, is converted into slow organ-scale movement (1–3). The ultra-slow tail movement described here may reflect an analogous mechanical principle in an animal body axis, despite the different molecular and material substrates involved. In the *Xenopus* tail, alternating left–right imbalance in growth, tissue compliance, or mechanical constraint could originate within the notochord–sheath system, through time-varying asymmetries in notochord cell expansion or in the formation and remodeling of the notochord sheath. Alternatively, it could be imposed by adjacent paraxial or somitic tissues, where segmentation-related gene-expression patterns can show consistent left–right phase differences in *Xenopus* (15).

The variable onset of this movement (Fig. 1G), together with the substantial elongation already present in late-onset embryos, suggests that it is not strictly required for axial elongation itself. Whether the movement subsequently participates in feedback regulation of tail morphogenesis, serves a role analogous to the exploratory function proposed for plant circumnutation, or instead represents an emergent readout of elongation mechanics remains to be determined.

In summary, we describe an exceptionally slow rhythmic movement in the elongating tail of *Xenopus* tadpoles. Its period scales with tail elongation rate and falls within the timescale range of slow movements in elongating plant structures. These findings raise the possibility that slow rhythmic motion is a recurrent dynamic feature of constrained elongation in living structures, suggesting a morphogenetic mode of animal movement that is closer to plant growth movements than to conventional behavior.

## Materials and Methods

*Xenopus laevis* embryos were maintained in 1/3× MMR containing pH-adjusted tricaine methanesulfonate (MS-222; final concentration, 0.3 mg/mL) during imaging. Long-term imaging from NF stage 22 was performed every 5 min using a modified document scanner, whereas temperature-controlled imaging from NF stage 35 was performed using a Raspberry Pi camera system at 13°C, 18°C, or 23°C. Experimental and analytical details are provided in Supplemental Information.

## Acknowledgments

We thank Asako Shindo, Shunya Kuroda, Koichi Fujimoto, Naoya Kamamoto, Tohya Suzuki, Yoshifumi Asakura, Tetsuya Nakamura, John B. Wallingford, and the members of the Shindo lab for valuable comments on this research. This study was supported by Grants-in-Aid for Scientific Research (23K19369 and 24H01488 to S.K.).

## Artificial intelligence disclosure

The authors used OpenAI ChatGPT (GPT-5.5 Thinking) to assist with English translation, language editing, formatting checks, and drafting Python analysis code. The authors reviewed, edited, and fact-checked all AI-assisted outputs and take full responsibility for the manuscript.

