## Supplemental Information for "Ultra-Slow Rhythmic Movement in the Elongating Tail of *Xenopus* Tadpoles"

**Author Contributions:** S.K. designed research with input from I.N.; I.N. and S.K. performed research; I.N. Y.A. and S.K. analyzed data; and S.K. wrote the paper with input from I.N.

**Competing Interest Statement:** The authors declare no competing interests.

**Keywords:** Morphogenesis, Tissue elongation, Rhythmic movement, *Xenopus*, Circumnutation

**This file includes:**

Extended methods
Video S1 to S3 descriptions

### Extended methods

#### *Xenopus* laevis tadpole preparation

Ovulation and mating of adult *Xenopus laevis* were induced by injection of human chorionic gonadotropin (Fuji Pharma, Japan) into the hindlimb base (100 U for males and 800 U for females). Eggs produced by natural mating after overnight incubation were collected and cultured in one-third strength Modified Marc's Ringer's (1/3× MMR) solution.

#### Time-lapse imaging of embryos

For all imaging experiments, embryos were maintained in 1/3× MMR (pH 7.4) containing pH-adjusted tricaine methanesulfonate (MS-222; Sigma-Aldrich, E10521; final concentration, 0.3 mg/mL).

For long-term imaging from NF stage 22 (Fig. 1A–G), embryos were imaged every 5 min using a document scanner (GT-X980, Epson). Stage 22 embryos were placed in a PDMS-coated dish, oriented dorsal side up, and immobilized with steel pins (Minutiens, 0.15 mm; BOHEMIA Insect Pins) placed bilaterally at the neck and anterior to the head. Dishes were covered with a custom-made cooled humid chamber to minimize medium evaporation and scanner-induced heating. Images were acquired from the ventral side using Epson Scan 2 controlled by PyAutoGUI. Only embryos that developed normally to at least NF stage 41 were included in the analysis.

For temperature-controlled imaging (Fig. 2), NF stage 35 embryos were imaged every 5 min using a Raspberry Pi High Quality Camera (RPI-SC1220, Raspberry Pi) equipped with a 16 mm C-mount telephoto lens (RPI-16MM-LENS, Raspberry Pi) and controlled by a Raspberry Pi 5. Embryos were placed in black PDMS-coated 35 mm dishes mounted on 65 mm stainless steel dishes, oriented dorsal side up, and immobilized with steel pins placed bilaterally at the neck. Dishes were incubated in a block incubator (BI-515S/BI-516S, ASTEC), and the medium temperature was maintained at 13°C, 18°C, or 23°C. Images were acquired from the dorsal side using custom Python code.

#### Quantification of tail-tip motion

Tail-tip motion was quantified using Fiji. Image stacks were first rotated such that the tail pointed leftward. Each frame was processed with Gaussian blur (sigma = 5) and binarized. The embryo contour was extracted using the Magic Wand tool, and the leftmost contour point, corresponding to the tail tip, was identified in each frame. The y-coordinate of this point was recorded as the lateral tail-tip position. For visualization, a 10-pixel-wide region of interest (ROI) centered on the tail-tip x-coordinate was extracted from each frame and assembled into a montage (Fig. 1D). The x-coordinate of the same point was used as a proxy for tail elongation in the temperature-rescaling analysis, with the sign inverted so that increasing values corresponded to elongation.

For the stage-aligned analysis shown in Fig. 1G, time points corresponding to NF stages 22 and 41 were identified morphologically in each embryo and used as developmental anchors. The intervening time axis was then linearly normalized to provide an approximate NF stage coordinate for comparing embryos. The resulting stage assignments were checked against embryo morphology in the raw images.

#### Detrending of tail-tip motion

The y-coordinate of the extracted tail-tip position was plotted against time. Long-term trends were removed by locally weighted scatterplot smoothing (LOESS) using smoothing parameters (frac) of 0.04–0.08 and 10 iterations. Detrended tail-tip motion was defined as the residual after subtraction of the fitted trend from the original trace.

Wavelet analysis

Detrended tail-tip motion was analyzed by continuous wavelet transform (CWT) using PyWavelets with a complex Morlet wavelet (cmor1.5-1.0) and a scale spacing of dj = 0.01. Wavelet power was defined as the squared magnitude of the wavelet coefficients. Scalograms were generated over a period range of 10–600 min (Fig. 1F). To determine a representative oscillation period, mean CWT power was evaluated for each candidate period using a band spanning ±50% of that period, and the period with the highest band-averaged power was defined as the peak period. The corresponding ±50% band was then extracted and displayed (Fig. 1G).

#### Fast Fourier transform (FFT)-based analysis

Detrended tail-tip motion was subjected to FFT-based spectral analysis using a custom Python script. After mean subtraction, the one-sided FFT amplitude spectrum was computed. The dominant frequency was defined as the non-zero frequency with the highest amplitude, and the oscillation period was calculated as its reciprocal (Fig. 2C and 2E).

#### Temperature rescaling

For temperature rescaling, the x-coordinate of the tail tip was used as a proxy for tail elongation (Fig. 2B). For each temperature condition, the temporal change in tail-tip x-coordinate was fitted by linear regression, and the time axis was rescaled according to the slope of the fitted line so that the fitted elongation rate matched that of the 23°C condition (Fig. 2D).

#### Data analysis and software

Image analysis was performed using Fiji and custom Python scripts. Detrending was performed by LOESS, and wavelet analysis was performed using PyWavelets. Statistical summaries and plots were prepared in GraphPad Prism and Microsoft Excel. Figures and movies were assembled using Affinity Designer and DaVinci Resolve, respectively.

#### Data and code availability

The custom code used for the analyses in this study is publicly available at GitHub (https://github.com/skato-lab/xenopus-tail-wagging-analysis) and archived at Zenodo (https://doi.org/10.5281/zenodo.19399806). Processed data supporting the findings of this study are available from the corresponding author upon reasonable request during peer review and will be deposited in Zenodo before journal publication. Raw image sequences are available from the corresponding author upon reasonable request.

#### Ethics statement

All animal experiments were approved by the Animal Care and Use Committee of The University of Osaka and were conducted in accordance with institutional guidelines.

### Video descriptions

**Video S1. Ultra-slow tail rhythmic movement begins during tail elongation**

Long-term ventral time-lapse imaging of a *Xenopus* embryo from St. 22 onward. The upper panel shows the whole embryo, and the lower panel shows a temporal montage of the tail-tip region indicated by the red band. Rhythmic left–right movement of the tail tip becomes apparent at around 20 h. Related to Fig. 1A–D.

**Video S2. Comparison of rhythmic tail movement periods across temperatures**

The left panel shows enlarged time-lapse views of the tail tip, and the right panel shows LOESS-detrended traces of tail-tip displacement over time. From top to bottom, the panels correspond to 13°C, 18°C, and 23°C. Related to Fig. 2A–C.

**Video S3. Comparison of rhythmic tail movement periods across temperatures after rescaling of time by tail elongation rate.**

The left panel shows enlarged time-lapse views of the tail tip after time rescaling by tail elongation rate, and the right panel shows LOESS-detrended traces of tail-tip displacement on the rescaled time axis. From top to bottom, the panels correspond to 13°C, 18°C, and 23°C. Related to Fig. 2D and E.
